# Human immunoglobulin from transchromosomic bovines hyperimmunized with SARS-CoV-2 spike antigen efficiently neutralizes viral variants

**DOI:** 10.1101/2021.02.06.430072

**Authors:** Zhuoming Liu, Hua Wu, Kristi A. Egland, Theron C. Gilliland, Matthew D. Dunn, Thomas C. Luke, Eddie J. Sullivan, William B. Klimstra, Christoph L. Bausch, Sean P. J. Whelan

## Abstract

Severe acute respiratory syndrome coronavirus 2 (SARS-CoV-2) variants with amino-acid substitutions and deletions in spike protein (S) can reduce the effectiveness of monoclonal antibodies (mAbs) and may compromise immunity induced by vaccines. We report a polyclonal, fully human, anti-SARS-CoV-2 immunoglobulin produced in transchromosomic bovines (Tc-hIgG-SARS-CoV-2) hyperimmunized with two doses of plasmid DNA encoding the SARS-CoV-2 Wuhan strain S gene, followed by repeated immunization with S protein purified from insect cells. The resulting Tc-hIgG-SARS-CoV-2, termed SAB-185, efficiently neutralizes SARS-CoV-2, and vesicular stomatitis virus (VSV) SARS-CoV-2 chimeras *in vitro*. Neutralization potency was retained for S variants including S477N, E484K, and N501Y, substitutions present in recent variants of concern. In contrast to the ease of selection of escape variants with mAbs and convalescent human plasma, we were unable to isolate VSV-SARS-CoV-2 mutants resistant to Tc-hIgG-SARS-CoV-2 neutralization. This fully human immunoglobulin that potently inhibits SARS-CoV-2 infection may provide an effective therapeutic to combat COVID-19.

## Introduction

Ending the global pandemic caused by severe acute respiratory syndrome coronavirus 2 (SARS-CoV-2) requires the deployment of multiple countermeasures including therapeutics and vaccines. Emergency use authorization (EUA) has already been granted by the United States Food and Drug Administration for vaccines, monoclonal antibodies (mAbs) and convalescent plasma (Baden et al., 2021; Bloch et al., 2020; Graham, 2020; Group et al., 2020; Halfmann et al., 2020; Krammer, 2020; Polack et al., 2020; Weinreich et al., 2020). A shared feature of those countermeasures are neutralizing antibodies that target the spike protein (S) of SARS-CoV-2. The S protein is responsible for engaging the viral receptor, angiotensin converting enzyme 2 (ACE2), and catalyzes fusion of the viral and host cell membranes initiating the process of infection (Letko et al., 2020). Genomic analysis of circulating SARS-CoV-2 variants has identified mutations in S including E484K that blunt the ability of multiple mAbs and convalescent plasma to neutralize virus in cell culture based assays (Allison J. Greaney, 2021; Baum et al., 2020; Greaney et al., 2020; Weisblum et al., 2020; Z Liu., 2021). Substitutions N501Y (B.1.1.7) and E484K-N501Y (B.1.351) are present among variants of concern which also possess other mutations in the spike gene and elsewhere in the viral genome (C Rees-Spear, 2021; Hou et al., 2020; Naveca F, 2021; Rambaut A, 2020; Tegally et al., 2020). To date, sera from vaccinated individuals neutralize the new variants in cell culture assays with some loss of potency. The impact of S mutations on the efficacy of therapeutic mAbs and convalescent plasma warrants the development of additional countermeasures (C K Wibmer, 2021; P Wang., 2021; Z Wang, 2021).

Genetically modified transchromosomic bovines (Tc-bovines) adaptively produce fully human polyclonal antibodies after exposure to environmental or vaccine antigens (Kuroiwa et al., 2009; Luke et al., 2018; Luke et al., 2016; Matsushita et al., 2014). After hyperimmunization, Tc-bovines produce high titer, fully human IgG (Tc-hIgG) that can be rapidly produced from their plasma (Kuroiwa et al., 2009; Luke et al., 2018; Luke et al., 2016; Matsushita et al., 2015). Tc-hIgGs have shown pre-clinical efficacy against Middle East Respiratory Syndrome Coronavirus (MERS-CoV), Hantaan, Ebola and Venezuelan Equine Encephalitis viruses among others (Casey C. Perley et al., 2020; Gardner et al., 2017; Luke et al., 2018; Luke et al., 2016). Previous randomized, double blind, Phase 1 or 1b clinical trials against MERS-CoV and mycoplasma hominis (ClinicalTrials.gov nos., NCT02788188 and NCT02508584 respectively) found Tc-hIgG to be safe, well tolerated and non-immunogenic (Beigel et al., 2018; Silver, et al., 2018).

Here we used a fully-human, polyclonal anti-SARS-CoV-2 immunoglobulin Tc-hIgG-SARS-CoV-2, termed SAB-185, produced from Tc bovines hyperimmunized with two doses of plasmid DNA encoding the SARS-CoV-2 Wuhan-Hu-1 strain Spike (S) gene (Wu et al., 2020), followed by repeated doses of recombinant S protein. Evaluation of SAB-185 in a Phase 1 (healthy adult) and Phase 1b (non-hospitalized SARS-CoV-2 infected) clinical trial (ClinicalTrials.gov nos., NCT04468958 and NCT0446917 respectively) revealed SAB-185 to be safe and well tolerated (manuscript in preparation). Distinct mutants to the Wuhan-Hu-1 virus strain, such as the D614G strain, and three newer mutants from the United Kingdom, South Africa and Brazil, have arisen (C K Wibmer, 2021; Hou et al., 2020; Naveca F, 2021; Rambaut A, 2020; Tegally et al., 2020). We therefore tested the ability of this Tc-hIgG-SARS-CoV-2 to neutralize SARS-CoV-2 (D614G variant) and several VSV-SARS-CoV-2 chimeric viruses *in vitro*. We demonstrate that SAB-185 retains potent neutralizing activity against the D614G, S477N, E484K, and N501Y S protein substitutions in cell culture based assays and were unable to isolate escape variants. This finding contrasts with the ability to isolate escape variants with convalescent human plasma and several neutralizing mAbs.

## Results

### Production of polyclonal human IgG against SARS-CoV-2 spike in transchromosomic bovines

To generate anti-SARS-CoV-2 polyclonal human immunoglobulin, we primed Tc-bovines with a DNA encoding the Wuhan-Hu-1 strain Spike (S) gene (Wu et al., 2020) for the first vaccination (V1) and the second vaccination (V2) at a 3-week interval, followed by 3 subsequent boosts with recombinant spike ectodomain produced and purified from insect cells for the third vaccination (V3) to the fifth vaccination (V5) at a 4-week interval (**Figure 1A**). Plasma was collected at 8, 11 and 14 days post V3 to V5 boosts and subjected to cGMP IgG purification to produce SAB-185 (**Figure 1A**). We selected multiple lots of SAB-185 for further analysis: Lot 1 obtained from V3 plasma, Lot 5 from V4 plasma, and Lot 6 from pooled V3, V4 and V5 plasma. The titers of SARS-CoV-2 spike specific IgG titers for each lot were evaluated by ELISA compared to human IgG purified Tc bovine pre-immune plasma as a negative control (**Figure 1B**). As expected, the neutralization titers increased with extent of immunization, with Lot 5 and Lot 6 having higher titer than Lot 1.

**Figure 1:**
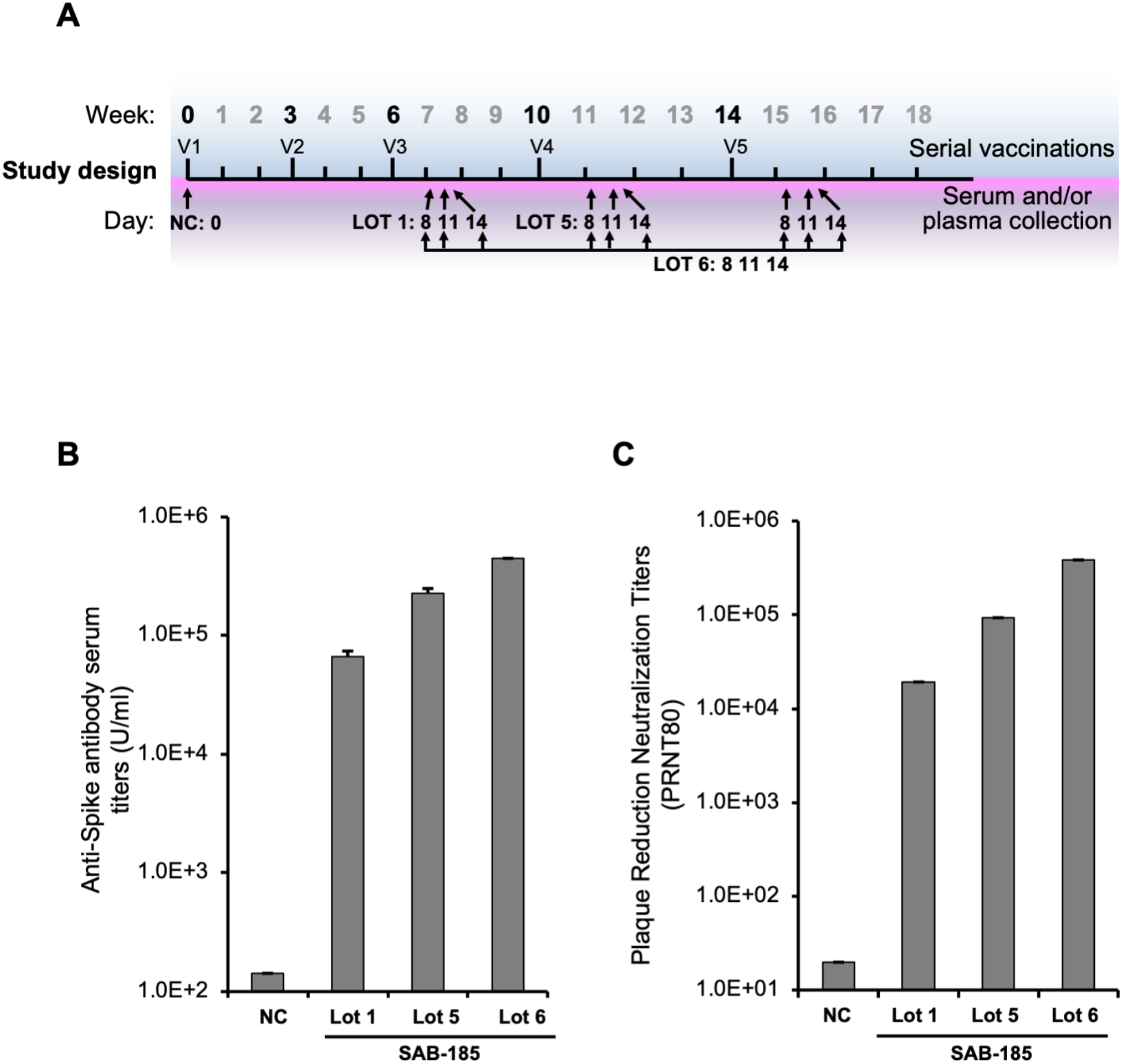
Generation of SARS-CoV-2 S SAB-185 polyclonal antibody. (**A**) Schedule of Tc bovine vaccinations V1 to V5, serum/plasma collection and plasma pooled for SAB-185 Lot 1, Lot 5 and Lot 6 purification as described in Materials and Methods. (**B**) Geometric mean SARS-CoV-2 spike ELISA antibody titers of SAB-185 Lot 1, 5 and 6 versus the negative control human IgG purified from Tc bovine pre-immune plasma. The negative control geometric mean titer (GMT) was calculated from six experiments. The Lot 1 GMT was from four experiments, Lot 5 GMT was from two experiments and Lot 6 was from a single experiment. (**C**) PRNT80 titers against the Munich SARS-CoV-2 P3 strain (Spike D614G) for a negative control pAb and SAB-185 Lot 1, Lot 5 and Lot 6. Concentrations of each pAb were normalized prior to serial 2-fold dilutions and 80% neutralization endpoints were calculated as described in Materials and Methods. The negative control endpoint was calculated from six replicate wells averaged at a 1:20 dilution of pAb. The Lot 1 endpoint was calculated from an average of two wells on a single plate (2 wells total) and Lot 5/Lot 6 were calculated from an average of two wells on three separate plates (6 wells total).

### Effect of Tc-hIgG-SARS-CoV-2 on neutralization of SARS-CoV-2

To examine whether this purified polyclonal human IgG could neutralize SARS-CoV-2, we performed a plaque reduction neutralization test assay using SARS-CoV-2 with the D614G (Munich strain) S substitution. The neutralization potency of Lot 1, 5 and 6 increased (**Figure 1C**). Similar results were obtained from a neutralization assay using a chimeric VSV-SARS-CoV-2 (Case et al., 2020) (**Figure 2A**). For VSV-SARS-CoV-2 we obtained IC_50_ of 4730 ng ml^−1^ with Lot 1, 650 ng ml^−1^ for Lot 5 and 212 ng ml^−1^ for Lot 6 corresponding to their levels of SARS-CoV-2 specific IgG. The IC_50_ values for virus neutralization compare favorably to those of potently neutralizing mAbs including 2H04 (**Figure 2A**).

**Figure 2.**
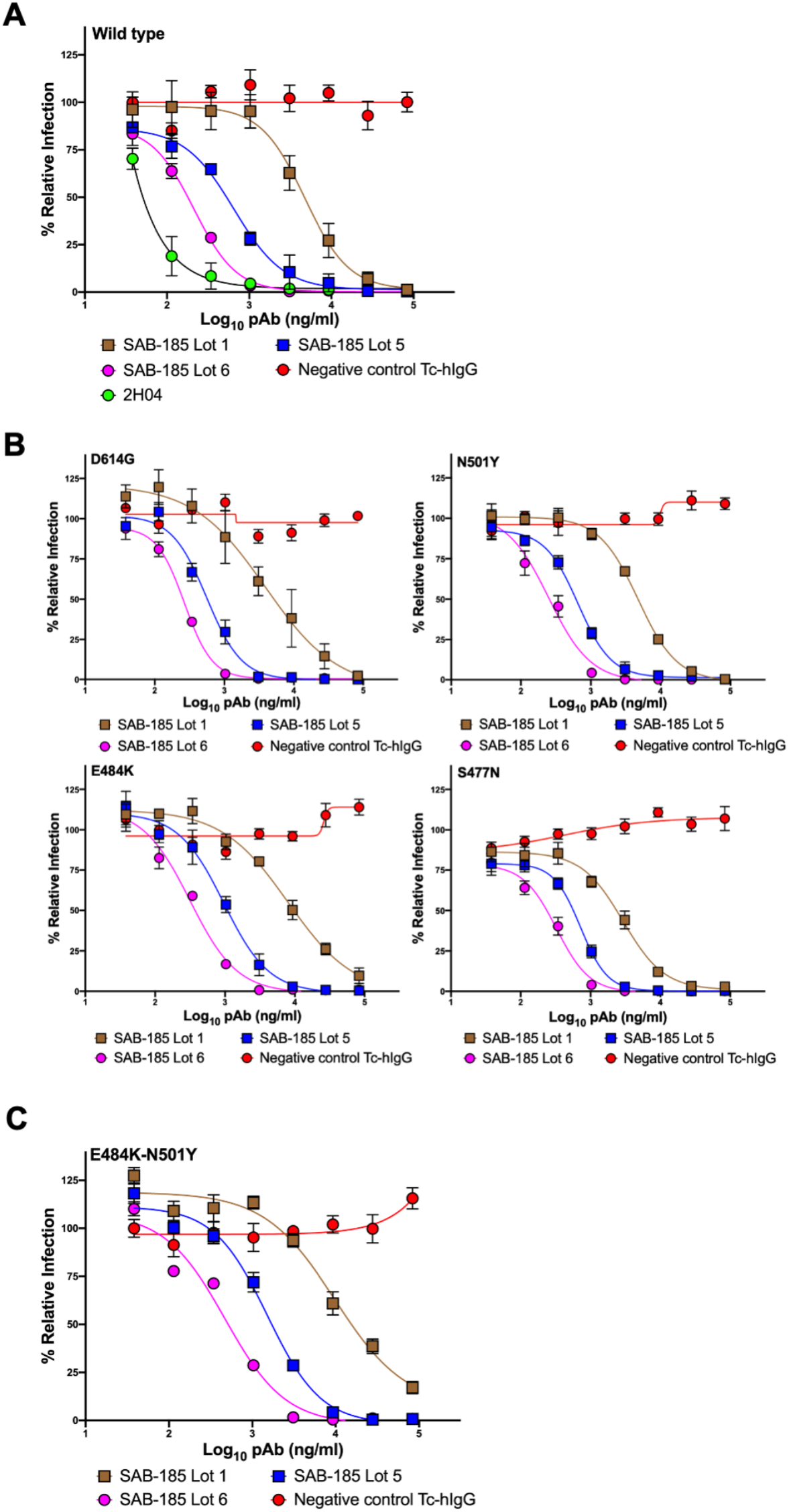
Neutralization of VSV-SARS-CoV-2 mutants by polyclonal antibody. **(A)** three SAB-185 pAbs were tested for neutralization of wild-type and (**B**) single amino acid substitution and (**C**) two amino acid substitution mutant VSV-SARS-CoV-2 (n = 4). Error bars represent the SEM. Data are representative of four independent experiments.

We previously selected >50 mutants in SARS-CoV-2 spike that exhibit resistance to specific monoclonal antibodies (Z Liu., 2021). Among those mutants were S477N and E484K which exhibit resistance to multiple mAbs and are present in emerging variants of concern. We also generated the dominant D614G, and the mouse adapted N501Y (Gu et al., 2020; Hou et al., 2020) and E484K-N501Yvariants. To determine whether the potency of SAB-185 was altered by any of these individual amino acid substitutions we performed neutralization assays with the corresponding chimeric VSV (**Figure 2B**). Each mutant exhibits a dose-dependent inhibition of infection by Lots 1, 5 and 6 of SAB-185, at levels that were similar to wild type S. We therefore tested whether the Tc-hIgG-SARS-CoV-2 also retains potency against a combination of substitutions E484K-N501Y in S. The resulting chimeric virus was also potently neutralized by SAB-185 with IC_50_ values of 9744 ng ml^−1^, 1508 ng ml^−1^ and 468 ng ml^−1^ for Lots 1, 5, and 6 (**Figure 2C**). This data demonstrates that SAB-185 retains neutralizing potency against multiple substitutions in S in chimeric VSV cell culture assays including several present in circulating human variants of SARS-CoV-2.

### Selections for resistance mutations using chimeric VSV-SARS-CoV-2

We and others have previously isolated VSV-SARS-CoV-2 S gene mutants by selection using mAbs and human convalescent serum that are resistant to neutralization (Baum et al., 2020; Weisblum et al., 2020; Z Liu., 2021). We therefore applied the same approach in an attempt to isolate variants resistant to SAB-185. In contrast to the ability to readily isolate mAb and convalescent serum escape mutants, we were unable to isolate mutants resistant to the human immunoglobulin SAB-185 from 3 successive attempts at sub-neutralizing concentrations of SAB-185 (**Figure 3**). Taken together, this analysis suggests that SAB-185 poses a significant barrier to immune escape.

**Figure 3.**
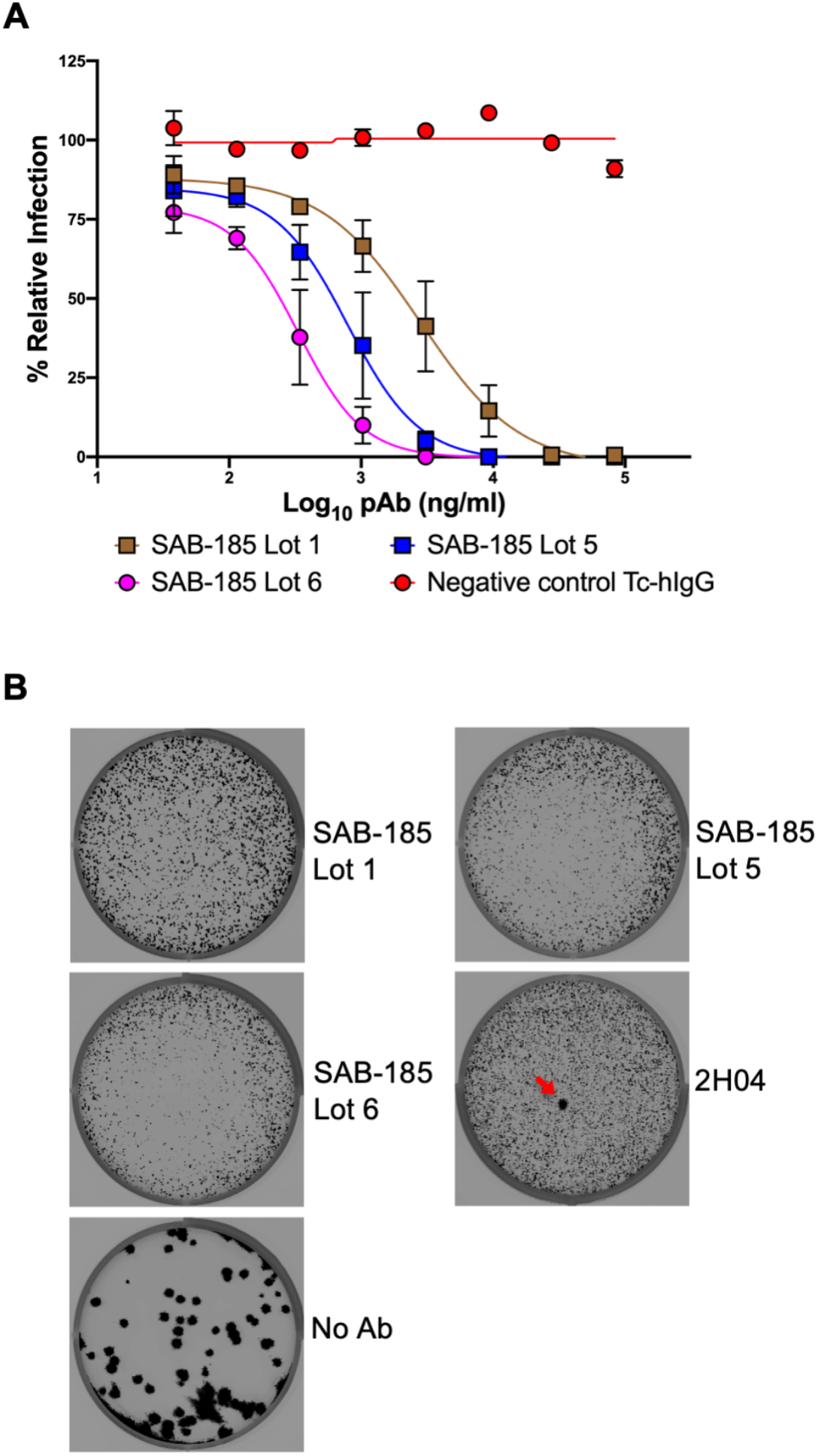
Selection of SAB-185 pAbs escape. (**A**) SAB-185 pAbs were tested for neutralizing activity against VSV-SARS-CoV-2 using an MOI of 1 to determine the concentration added into overlay. Error bars represent the SEM. Data are representative of four independent experiments. (B) Plaque assays were performed to isolate the VSV-SARS-CoV-2-S escape mutant on Vero E6 TMPRSS2 cells in the present of the indicated pAb in the overlay. The concentration of SAB-185 pAbs added in the overlay completely inhibited viral infection (**See Figure 3A**). Representative images of eight independent experiments are shown.

## Discussion

We report a potently neutralizing polyclonal human IgG produced in transchromosomic bovines that effectively neutralizes SARS-CoV-2 and VSV-SARS-CoV-2 chimeras that contain one or two S mutations present in circulating variants of concern (Greaney et al., 2020; Li et al., 2020; Weisblum et al., 2020; Z Liu., 2021). This finding contrasts with earlier work which shows that immune sera obtained from patients that have recovered from COVID-19 as well as several mAbs show significant reductions in neutralization potency in cell culture assays particularly against S substitutions E484K and S477N. The potency seen for SAB-185 with the combination of E484K and N501Y likely reflects the fact that the hyperimmune IgG SAB-185 recognizes a broader swath of epitopes in S such that single substitutions are unlikely to escape neutralization. As a result, large scale antigenic variation that results in the appearance of new serotypes may be required to evade neutralization by such a polyclonal human IgG.

The emergence of new SARS-CoV-2 variants in humans and other species will continue to pose a threat to human health particularly if sufficient antigenic variation is achieved to blunt vaccine induced immunity (Dinnon et al., 2020; Gu et al., 2020; Halfmann et al., 2020; Oude Munnink et al., 2020; Shi et al., 2020). It seems likely, however, that pre-existing immunity developed as a consequence of natural infection or through vaccination may offer some protection from disease. The present study evaluated the ability of the polyclonal IgG SAB-185 to neutralize virus in cell culture assays, and how this impacts disease outcome will require studies in vivo. In addition to neutralizing antibodies, non-neutralizing antibodies can offer protection of animals from infection by SARS-CoV-2 in experimental models of disease (Winkler et al., 2020). The extent to which natural infection or vaccination induces non-neutralizing but protective antibodies is uncertain. Both vaccination and natural infection will also stimulate a T-cell response that also contributes to immunity (Lu et al., 2018).

Production of human IgG by transchromosomic cattle has been described for multiple emerging viruses, including Ebola, Hantaan, and MERS CoV (Casey C. Perley et al., 2020; Luke et al., 2018; Luke et al., 2016), demonstrating that it provides a versatile platform for development of immunotherapeutics. Identification of the key antigenic determinants is critical, but in the case of enveloped viruses their surface proteins are typically effective targets. Using nucleic acid as the immunogen also limits the need for production of recombinant protein or vectored vaccine approaches that may pose unexpected challenges toward developing human IgG. Here we used recombinant S produced in insect cells to boost the immune response following priming with nucleic acid. Although there are differences in glycosylation in insect cells, the recombinant protein effectively boosted neutralizing IgG titers as demonstrated by the increased potency of Lot 6 and 5 over Lot 1. In phase 1 clinical trials, SAB-185 was shown to be safe and well tolerated and given its potent neutralization efficacy in vitro, use of SAB-185 may provide a useful addition to the evolving repertoire of countermeasures against COVID-19.

## ACKNOWLEDGEMENTS

We thank Ma. Xenia G. Ilagan in automated microscope core facility, Spencer Stumpf for excellent technical assistance and Ali H. Ellebedy for generously providing 2H04.

## AUTHOR CONTRIBUTIONS

Z.L designed and performed the VSV experiments. W.H and K.A.E produced the purified SARS-CoV-2 Spike pDNA and protein antigens. W.H immunized the Tc-bovines. C.L.B generated and purified SAB-185. T.C.G and M.D.D performed the neutralization assay at the Biosafety Level 3 facility. Z.L, H.W, T.C.L, W.B.K, E.J.S and S.P.J.W analyzed the data. Z.L, H.W, T.C.L, W.B.K, and S.P.J.W wrote the initial draft, with the other authors providing editing comments.

## EXPERIMENTAL MODEL AND SUBJECT DETAILS

### Cells

Cells were cultured in humidified incubators at 34° or 37°C and 5% CO_2_ in the indicated media. Vero CCL81, Vero E6 and Vero E6-TMPRSS2 were maintained in DMEM (Corning or VWR) supplemented with glucose, L-glutamine, sodium pyruvate, and 10% fetal bovine serum (FBS). MA104 cells were propagated in Medium 199 (Gibco) containing 10% FBS. Vero E6-TMPRSS2 cells were generated using a lentivirus vector described as previously (Case et al., 2020).

### VSV-SARS-CoV-2 mutants

VSV-SARS-CoV-2 was described as previously (Case et al., 2020). S477N and E484K were escape mutants isolated from mAbs described as previously (Z Liu., 2021). N501Y and D614G were constructed using SARS-CoV-2 Wuhan-Hu-1 spike with substitution at N501 or D614 site respectively and rescued by using reverse genetic system. E484K-N501Y were escape mutants isolated from 2B04 using N501Y virus. Virus were then plaque purified and the mutations were identified by Sanger sequencing (GENEWIZ). Viral stocks were amplified on MA104 cells at an MOI of 0.01 in Medium 199 containing 2% FBS and 20 mM HEPES pH 7.7 (Millipore Sigma) at 34°C. Viral supernatants were harvested upon extensive cytopathic effect and clarified of cell debris by centrifugation at 1,000 x g for 5 min. Aliquots were maintained at −80°C.

## METHOD DETAILS

### Investigational new animal drug (INAD) and ethics statement

SAB Biotherapeutics has an Investigational New Animal Drug (INAD) file (#I-011204) with FDA Center for Veterinary Medicine (CVM) on the complete genetic engineering of the Transchromosomic (Tc) bovine and the production of fully human antibody in the animals. SAB uses Association for Assessment and Accreditation of Laboratory Animal Care International (AAALAC) for accreditation of its animal care and use programs. The animal protocols contained in the study were approved by SAB Biotherapeutic Institutional Animal Care and Use Committee (IACUC).

### Tc bovines

Tc bovines were produced as previously described(Kuroiwa et al., 2009; Luke et al., 2016). The Tc bovines used in this study are homozygous for either triple or quadruple knock-outs in the endogenous bovine immunoglobulin genes (*IGHM* − / − *IGHML1* − / − *IGL* − / − or *IGHM* − / − *IGHML1* − / − *IGL − / − /IGK− / −*) and carry a human artificial chromosome (HAC) vector labeled as isKcHACD with an IgG1 production bias. This HAC vector consists of human chromosome 14 fragment, which contains the entire human immunoglobulin heavy chain locus except that the IGHM constant region remains bovine and the key regulatory sequences were bovinized; and human chromosome 2 fragment, which contains the entire human immunoglobulin k light chain locus (Kuroiwa et al., 2009).

### Generation of pCAGGS and Spike fusion protein for Tc Bovine immunization

A plasmid encoding SARS-CoV-2 Spike protein and a purified Spike fusion protein were generated for use as antigens to stimulate production of human polyclonal antibodies using the Tc bovine platform. The pCAGGS expression plasmid, provided by Florian Krammer from Icahn School of Medicine at Mount Sinai, contains the SARS-CoV-2 Wuhan-Hu-1 sequence encoding the full-length Spike protein. Expression of the *spike* gene is regulated by the CMV promoter. Plasmid DNA was generated using the Qiagen EndoFree Plasmid Giga Kit. To produce purified Spike protein, the Bac-to-Bac Baculovirus Expression System (Gibco, Gaithersburg, MD) was used to make the antigen in the insect cell line, ExpiSf9, according to manufacturer’s instructions. Briefly, the SARS-CoV-2 Wuhan-Hu-1 sequence encoding the ectodomain of Spike (amino acids 1-1213) was cloned into pFastBAC1 3’ of the polyhedrin promoter with the following modifications. The furin cleavage site, RRAR (residues 682-685), was changed to GSAS. Substitutions K986P and V987P were added to stabilize the C-terminal S2 fusion machinery. The sequence encoding the 27 amino acid T4 foldon trimerization domain(Guthe et al., 2004) was cloned 3’ of the ectodomain. Linker sequences were added 5’ and 3’ of the encoded TEV protease recognition site, ENLYFQG, followed by six histidines. The sequence was codon optimized for insect expression systems, and the final plasmid is referred to as pSAB-uSpike. The pSAB-uSpike was transformed into competent DH10Bac *E. coli* containing a bacmid and helper vector to generate a recombinant bacmid. The purified uSpike bacmid was transfected into ExpiSf9 cells to produce P0 recombinant baculovirus stock. After viral amplification and titer determination, ExpiSf9 cells were infected with an MOI of 5. ExpiSF9 cells were grown at 27°C in a non-humidified and non-CO_2_ atmosphere incubator on an orbital shaker platform set at 125 ±5 rpm. After 96-120 hours, supernatants were harvested, and the purification procedures were performed at 4°C. His-tagged uSpike was purified using Ni-NTA agarose (Qiagen, Germantown, MD) in a gravity flow column. After washing and elution from the Ni-column, the imidazole was removed by Amicon centrifugal filters (Sigma Aldrich, St. Louis, MO). The his-tag was cleaved using AcTEV protease according to manufacturer’s instructions (Invitrogen, Carlsbad, CA), and the his-tagged protease and noncleaved uSpike protein were removed by binding to Ni-NTA resin. The purified uSpike antigen was buffer exchanged into PBS, and the protein concentration was determined by BCA assay.

### Tc bovine immunization and plasma collection

A group of Tc bovines were primed with SARS CoV2 12 mg pDNA along with SAB’s proprietary adjuvant formulation (SAB-adj-1) for the first vaccination (V1) and the second vaccination (V2) at three weeks interval, and then boosted with 2 to 5mg recombinant ectodomain of spike protein formulated with SAB-adi-1 for third vaccination (V3) to Fifth vaccination (V5) at 4-week intervals. pDNA was delivered by by using the PharmaJet Stratis^®^ IM injection device as previously described (Hooper et al., 2014). Up to 2.1% of body weight of hyperimmune plasma per animal was collected from immunized Tc bovines on days 8, 11 and 14 after each vaccination starting fromV3–V5. Plasma was collected using an automated plasmapheresis system (Baxter Healthcare, Autopheresis C Model 200). Plasma samples were stored frozen at −20°C until purifications were performed.

### SAB-185 cGMP Purification

After quality control testing, the qualified Tc bovine plasma was thawed, pooled, fractionated by caprylic acid (CA), and clarified by depth filtration in the presence of Celpure P1000 filter aid. The clarified sample containing Tc bovine-derived human IgG is further purified by affinity chromatography, first using an anti-human IgG kappa light chain specific column, KappaSelect (GE Healthcare Life Sciences) to capture hIgG and remove residual non-hIgG and bovine plasma proteins (BPP) followed by a low pH treatment, and, second, by passing through an anti-bovine IgG heavy chain specific affinity column, Capto HC15 (GE Healthcare Life Sciences). To further remove residual IgG that contains bovine heavy chain, the human IgG fraction was then subjected to a Q Sepharose chromatography polishing step to further reduce impurities, nanofiltration, final buffer exchange, concentration and sterile filtration. Finally, the SAB-185 product was terminally filtered and filled into vials. The product protein concentration was determined by A280 using the Unchained Labs Big Lunatic. The product SAB-185 was in a buffer at a pH of 5.5 consisting of 10 mM of glutamic acid monosodium salt, 262 mM of d-sorbitol, and 0.05 mg/mL of Tween 80. After quality control testing and quality assurance review, SAB-185 was released for preclinical and/or clinical studies. SAB-185 Lot 1 and Lot 5 were purified from pooled V3 plasma and V4 plasma, respectively. SAB-185 Lot 6 was purified from pooled V3, V4 and V5 plasma. The protein concentration for each lot of SAB-185 was as follows: Lot 1 at 75.48mg ml^−1^, Lot 5 at 73.01mg ml^−1^ and Lot 6 at 75.69mg ml^−1^.

### SARS CoV-2 spike-protein-specific human IgG ELISA

In this ELISA assays, recombinant ectodomain of SARS CoV-2 spike protein produced and purified from 293 cells was used as the coating antigen. Determination of SARS CoV-2 spike protein-specific human IgG antibody titers was performed in Maxisorp Immuno 96-well ELISA plates (Thermo Scientific) coated overnight at 4°C with 100 μl/well of 2 μg ml^−1^ recombinant SARS CoV-2 spike protein produced and purified from 293 cells in PBS. Plates were washed with PBST (PBS with 0.05% Tween 20) and blocked at RT for 1 hour with 1% bovine serum albumin in PBS. After washing with PBST, SAB-185 Lot1, 5 and 6 were serially diluted in PBST, added to the plates, and incubated for 1 hour at RT. Following washing with PBST, diluted goat anti-human IgG-Fc conjugated with horseradish peroxidase (HRP) (Bethyl) was added to plates and incubated for 1 hour at RT. After final washing with PBST, the bound anti-spike antibodies were detected colorimetrically by using the 3, 3,’ 5, 5’-tetramethylbenzidine (TMB) substrate kit (SeraCare). Absorbance was read in a microplate reader at 450 nm. The titer (units/ml) is defined as the reciprocal of the highest dilution of SAB-185 resulting in an optical density at 450 nm (OD450) reading that was 2.5-fold higher than blank.

### Plaque Reduction Neutralization Assay and plaque assay

Virus production and neutralization of a low passage strain of SARS-CoV-2 were as described as previously (Klimstra WB, 2020). Virus growth and assays were performed at Biosafety Level (BSL) 3 BSL-3 conditions in the Regional Biocontainment Laboratory (RBL) in the Center from Vaccine Research, at the University of Pittsburgh. Briefly, approximately 100 plaque forming units of SARS-CoV-2 were mixed with serial 10-fold dilutions of SAb-185, beginning with a 1:20 dilution, and incubated at 37°C for 1 hour followed by infection of two wells of a six-well plate of Vero E6 cells. Infection proceeded for 1 hour at 37°C followed by overlay with virus growth medium containing 0.1% (w/v) immunodiffusion agarose (MP Biomedicals) and incubation was continued for 96 hours. Plates were fixed with formaldehyde (37% (w/v) formaldehyde stabilized with 10-15% (v/v) methanol; Fisher Scientific) for 15 minutes at room temperature. Agarose and fixative were discarded and 1 ml/well 1% (w/v) crystal violet in 10% (v/v) methanol (both Fisher Scientific) was added followed by enumeration of plaques. Neutralization assays were performed in triplicate (6 total wells) and values averaged and then 1/10 of the reciprocal of the highest dilution at which 80% of control plate plaques were neutralized was taken as the PRNT_80_ value. VSV-SARS-CoV-2 plaque assays were performed on Vero and Vero E6-TMPRSS2 cells. Briefly, cells were seeded into 6 well plates for overnight. Virus was serially diluted using DMEM and cells were infected at 37°C for 1 h. Cells were cultured with an agarose overlay in the presence of Ab or absence of Ab at 34°C for 2 days. The concentration of SAB-185 pAbs added in the overlay completely inhibited viral infection. Plates were scanned on a biomolecular imager and expression of eGFP is show at 48 hours post-infection.

### Neutralization assays using a recombinant VSV-SARS-CoV-2 and mutants

Briefly, the initial dilution of started at 83 ug mL^−1^ and was three-fold serially diluted in 96-well plates over eight dilutions. Indicated dilutions of SAB-185 pAbs were incubated with 10^2^ PFU of VSV-SARS-CoV-2 and mutants for 1 h at 37 °C. SAB-185 pAb-virus complexes then were added to Vero E6 cells in 96-well plates and incubated at 37 °C for 7.5 h. Cells were fixed at room temperature in 2% formaldehyde containing 10 μg/mL of Hoechst 33342 nuclear stain for 45 min. Fixative was replaced with PBS prior to imaging. Images were acquired using an In Cell 2000 Analyzer automated microscope (GE Healthcare) in both the DAPI and FITC channels to visualize nuclei and infected cells (×4 objective, 4 fields per well). Images were analyzed using the Multi Target Analysis Module of the In Cell Analyzer 1000 Workstation Software (GE Healthcare). GFP-positive cells were identified using the top hat segmentation method and counted within the InCell Workstation software.

## QUANTIFICATION AND STATISTICAL ANALYSIS

All statistical tests were performed as described in the indicated figure legends. Non-linear regression (curve fit) was performed for Fig 2 and 3A using Prism 9.0. The number of independent experiments used are indicated in the relevant Figure legends.

